# Understanding Structural Mechanics of Ligated DNA Crystals via Molecular Dynamics Simulation

**DOI:** 10.1101/2025.07.17.665441

**Authors:** Yoo Hyun Kim, Anirudh S. Madhvacharyula, Ruixin Li, Alexander A. Swett, Seongmin Seo, Emile J. Batchelder-Schwab, Naseem Siraj, Chengde Mao, Jong Hyun Choi

## Abstract

DNA self-assembly is a highly programmable method that can construct arbitrary architectures based on sequence complementarity. Among various constructs, DNA crystals are macroscopic crystalline materials formed by assembling motifs via sticky end association. Due to their high structural integrity and size ranging from tens to hundreds of micrometers, DNA crystals offer unique opportunities to study structural properties and deformation behaviors of DNA assemblies. For example, enzymatic ligation of sticky ends can selectively seal nicks resulting in more robust structures with enhanced mechanical properties. However, the research efforts have been mostly on experiments such as different motif designs, structural optimization, or new synthesis methods, while their mechanics are not fully understood. The complex properties of DNA crystals are difficult to study via experiment alone, and numerical simulation can complement and inform the experiment. Coarse-grained molecular dynamics (MD) simulation is a powerful tool that can probe the mechanics of DNA assemblies. Here, we investigate DNA crystals made of four different motif lengths with various ligation patterns (full ligation, major directions, connectors, and in-plane) using oxDNA, an open-source, coarse-grained MD platform. We find that several distinct deformation stages emerge in response to mechanical loading and that the number and the location of the ligated nucleotides can significantly modulate structural behaviors. These findings should be useful for predicting crystal properties and thus improving the design.

## INTRODUCTION

DNA self-assembly is a powerful bottom-up approach that uses the Watson-Crick base pairing to construct complex structures with exceptional precision and reproducibility.^1^ Several strategies has been developed, including tile-based assembly^2, 3^, scaffolded origami^4-6^, and DNA bricks^7, 8^. In tile-based assembly, several DNA oligonucleotides form a motif which can assemble into 2D or 3D architectures via sticky end cohesion, often exploiting symmetry^9-11^. The DNA origami technique uses a long single-stranded (ss) DNA scaffold (e.g., 7249-base-long M13mp18) that is folded into an in-silico design stabilized by a set of staple oligos^4^. Lastly, then DNA-bricks approach uses short strands with unique sequences to assemble into arbitrary shapes via a one-step annealing process^3^. These methods have created static and dynamic DNA assemblies which can be used in a variety of applications such as biosensing^12-14^, drug delivery^15-17^, nanomachines^18-20^, and DNA computing^21, 22^. DNA assemblies are mostly in nanoscale dimensions on the order of 100 nm or less. There is a growing interest in scalable assembly methods toward large-scale DNA constructs.

DNA crystals are unique 3D crystalline structures which form by arranging DNA units in periodic patterns and connecting them via sticky end association. DNA units range from a small multi-strand motifs to DNA origami with a variety of geometries including triangle^23-25^, square^26, 27^, hexagonal^28^, and octahedral geometries^29^. This hierarchical approach can create unique 3D crystals with sizes exceeding 100 µm. In crystal construction, sticky end design and flexibility of the DNA units are important. The motifs typically have short overhangs (2 to 3 nucleotides long), compared to other constructs such as DNA bricks^7^ or reconfigurable structures^30-32^. While longer sticky ends can provide stronger cohesion and may lead to more resilient structures, they tend to form smaller crystals because of kinetic trapping of assembly defects. Shorter sticky ends allow more reversible binding and robust error correction process which results in large crystals with high resolution^33^. The flexibility/stiffness of the assembling units is also important, since a balance is needed between the unit design and its flexibility. While numerous advances have been made in the design and optimization of DNA structures, mechanical study has been limited to DNA duplexes^34-36^ or simple nanostructures^37-39^ due to the difficulties in force application and deformation measurements. Micron-sized DNA crystals can serve as an ideal platform for studying mechanical behavior of complex DNA structures, facilitating experiments.

Despite the potential as a versatile platform, there are also technical challenges in DNA crystals such as chemical stability and mechanical strength. The native crystals are fragile both chemically and mechanically. They are not stable at typical conditions for DNA self-assembly such as 12.5 mM Mg^2+^; instead, high ionic strength is required (e.g., 50 mM Mg^2+^). Their extreme sensitivity to temperature or dehydration also restrict their usage in applications^40^. Post-assembly modification such as silicification have shown possibilities of enhancing mechanical stability^41-43^ where the Young’s modulus of the DNA crystal was enhanced by three orders of magnitude^44^. Facile approaches also have been introduced to engineer crystals robust and resilient, including triplex formation^45^ and ligation of the sticky ends^40^. Ligase concentration and ligation time can be varied to modulate chemical and structural stability, thereby forming complex structures like York-shells and Matryoshka-dolls^40^. Ligation pattern and motif size were found to be among the key parameters that govern the chemical and mechanical properties of DNA crystals. However, their effects on material responses remain elusive, and thus, a better understanding will help develop design rules and improve hierarchical assembly methods for macroscopic crystals from DNA.

In this work, we investigate structural properties and mechanical behaviors of DNA crystals from tensegrity triangle motifs and elucidate the effects of ligation patterns and motif lengths under external loads using oxDNA, an open-source MD platform based on coarse-grained models. In a previous report, we showed that it is possible to study DNA crystals using a segment of moti assemblies with periodicity in oxDNA platform^46^. Building upon the earlier approach, we have examined four different ligation patterns: ligations along the major directions, connectors, and in-plane as well as full ligation. Motif lengths are varied from 2- to 8-turn-long. We observe stark differences as well as similarities in deformation behaviors in response to mechanical loading. These findings provide insights into fundamental structure-mechanical property relationships and will ultimately help improve the design of DNA assemblies.

## METHODS

### Motif Design

A tensegrity triangle motif is used for constructing 3D DNA crystal structures, which consists of seven strands. Three of those strands, in purple color in Fig. 1a-b, are oriented in three distinct (major) directions along x, y, and z axes and form the triangular framework. One center strand (cyan) holds the three strands together. Lastly, three connector strands (orange) have unpaired 2 or 3 nucleotides which are used as sticky ends and bind with purple strands from another motif. The association via sticky ends thus links motifs together into a 3D crystal^23^. The motif design is symmetrical, meaning that the DNA sequences within each strand type are identical. However, the edge length of each motif varies. In this work, four different edge lengths are introduced: two-, four-, six-, and eight-turn motifs (based on 10.5 base-pair/turn), termed 2T, 4T, 6T, and 8T, respectively. The corresponding sequences of each design were used in previous studies^23, 40^ and are provided in Supplementary Table S1.

**Figure 1.**
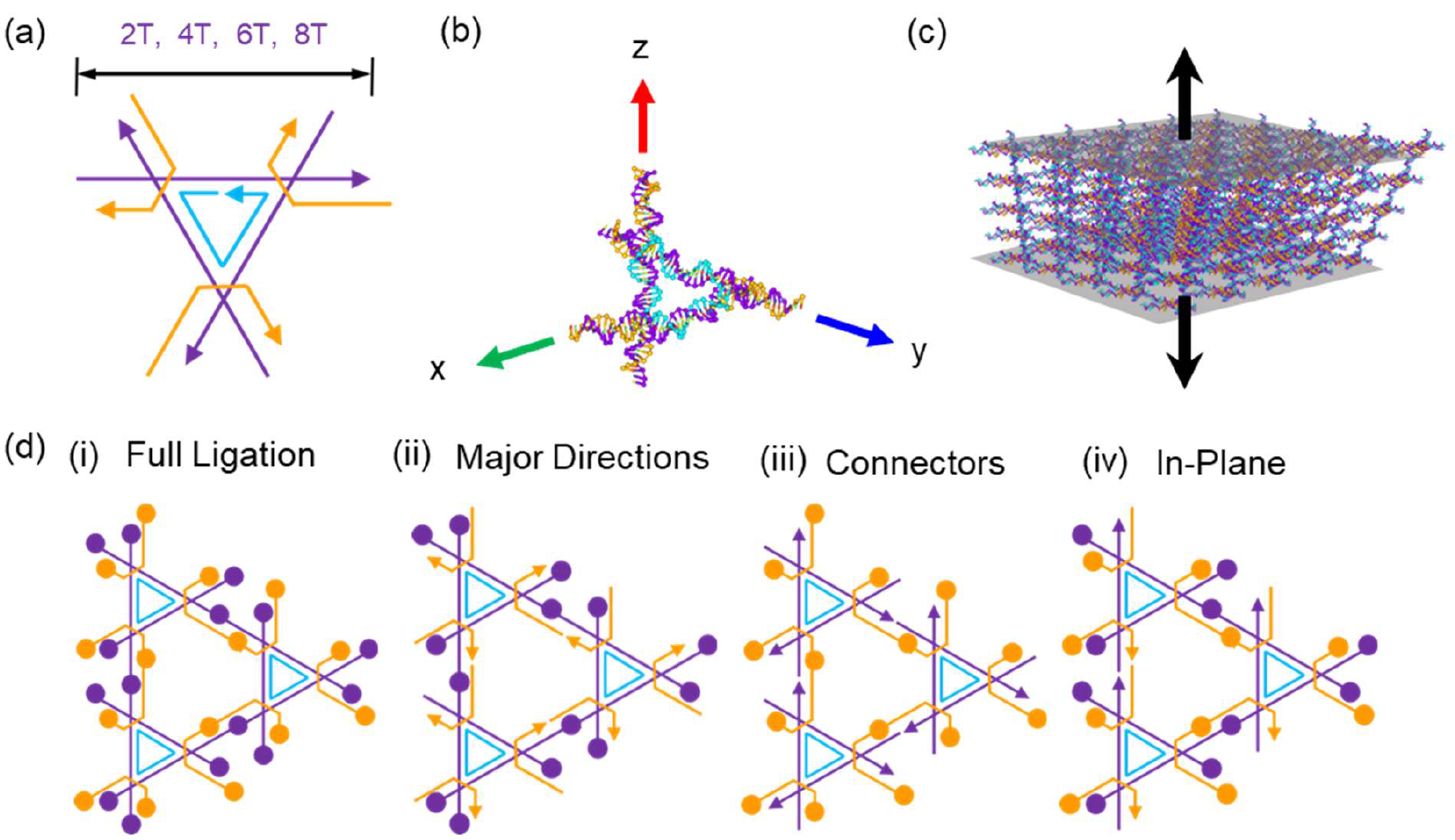
(a) Schematic of a single tensegrity triangle motif used for constructing DNA crystals. Each motif consists of seven strands, 1 cyan, 3 orange, and 3 purple strands. The length of the purple strands is varied for different numbers of helical turns; 2T, 4T, 6T, and 8T denote 2, 4, 6, and 8 helical turns, respectively. (b) A molecular model of the triangle motif. Three major directions are indicated with red, green, and blue arrows, where x and y axes are in-plane directions and z axis shown in red represent both height and loading direction. (c) Illustration of tensile force application on a crystal domain made of 5×5×5 motifs (a total of 125 units). The grey shaded regions represent the force planes, each containing 25 nucleotides from purple strands. The top plane is pulled upward while the bottom plane is fixed. The black arrows indicate the direction of the force applied on each plane. (d) Top view of three neighboring motifs, illustrating four different ligation patterns. The circles represent the ligation sites on purple and orange strands. Cyan strands are not marked for ligation because they are ligated in all cases.

### MD Simulation via oxDNA

MD methods simulate the movement of atoms and molecules over time by solving Newton’s equation of motion. While all-atom models can yield detailed results, it would take a long period of time, particularly for large structures^47, 48^. In coarse-grained models, a group of atoms are simplified, thus minimizing computing time. Coarse-grained MD approach has proven to be effective estimating equilibrium conformations and structural properties of large DNA constructs such as an origami structure^49, 50^. In this work, we used oxDNA,^51-53^ an open-source platform based on coarse-grained models, which allows for applications of external mechanical loading.

For all motif designs, the motif file was initially created using caDNAno2^54^ and converted into topology and configuration files compatible with the oxDNA simulation. Topology files include the sequence information of each strand. The motif was then duplicated into a total of 125 units, 5 times in each direction, forming a 5□5□5 domain by modifying the topology and configuration files. While a larger domain could potentially yield more accurate results, a balance between computational efficiency and accuracy is necessary. We found that the 5□5□5 crystal was sufficiently large to capture the deformation behavior under tensile loading while remaining feasible for simulation.

The domain structure was relaxed to reach an equilibrium state by running the simulation for 10 million steps. During the relaxation process, 25 nucleotides (one nucleotide from each motif in the bottom layer) as illustrated by the grey shaded region in Fig 1c, were fixed in place using a set of force planes. The nucleotides were positioned between two repulsion force planes that applied opposite forces, keeping the nucleotides at the center of the two planes^55^. After relaxation, a tensile force was exerted on the top plane (grey shade on top in Fig. 1c), which consisted of the top 25 nucleotides using a set of repulsion planes as well. The loading planes were moved upward, away from the bottom plane, and the force constant was increased to simulate the deformation process. To maximize the tension, the nucleotides on the top and bottom planes were selected from the purple strands that vertically pass through all five layers of planes. To securely fix the bottom plane, the stiffness of the bottom force planes was constantly maintained 100 times greater than that of the top force planes. For each data point (i.e., each conformation), the simulation was performed for 2 million steps with the temperature set at T = 300 K and the salt concentration at [Na^+^] = 0.5 M, to ensure that the structure reaches an equilibrium state. The equilibrium is defined as when the height of the structure has a relative fluctuation of less than 3%.

### Ligation Patterns

The crystal domains were constructed with four distinct ligation patterns to investigate the impact of ligation on structural properties and deformation behaviors. The ligations for simulation were performed using oxView^56, 57^ by selecting the 3’ ends of the ligating strand and 5’ ends of the adjacent strand. Upon ligation, it joins the two separate strands into a single continuous strand. First, we ligate both purple and orange strands in all directions, which we term ‘full ligation’ (Figure 1d(i)). When only purple strands are ligated, three major-directions are seamlessly connected. Thus, this ligation pattern is called ‘major directions’ (Figure 1d(ii)). We also examine another pattern where the orange connector strands are ligated (Figure 1b(iii)). Lastly, ligation is applied in x and y directions (Figure 1d(iv)), which makes the strands in-plane ligated while the z-direction (height; loading direction) remains native. Note that the cyan strands were ligated in all cases.

### Computational Analysis

The simulation results were analyzed after reaching equilibrium by obtaining position data of the nucleotides. The extension of the crystal domain was calculated by subtracting the average height of the bottom plane from that of the top plane. The magnitude of the applied force was computed as the product of stiffness and the distance between the top nucleotide plane and the force plane. The edge length of the cross section was obtained by calculating the vector distance between two opposite ends along each edge. The cross-sectional area was then determined by computing the magnitude of the cross product of the two edge vectors, representing the area of the parallelogram defined by the edges.

The data of hydrogen binding energy was collected for the calculation of base pairing ratio of DNA duplex. To determine the state of the base paring, a threshold value was defined as two standard deviations above the average hydrogen bond energy at the initial state, when no force is exerted and all the nucleotides are bound. This resulted in about -1.3 × 10^−20^ J. With this threshold value, the total number of base-paired nucleotides was counted for each simulation output, and the base paring ratio was calculated by dividing the number of hybridized base pairs by the total number of base pairs.

## RESULTS AND DISCUSSION

### Deformation Behavior of 4T Crystals Under Different Ligation Patterns

Our previous report investigated the mechanical behavior of a fully ligated 4T DNA crystal using both MD simulations and AFM nanoindentation^46^. The Young’s modulus of the structure was reported to be approximately 1 MPa. The simulation results were also compared with other experimental studies in which λ-DNA was stretched using optical tweezers. The comparison revealed similar deformation behavior, supporting the validity of the simulation model. Some differences were also noted, such as abrupt changes in double-stranded DNA (dsDNA) stretching and strain range across deformation stages. These findings highlight that while the mechanical behavior of a DNA crystal is strongly influenced by their components, additional factors such as motif design and ligation patterns must be considered to fully understand the deformation behaviors. To this end, various ligation patterns and motif lengths were tested under tensile loading to investigate the structural mechanics of DNA crystals in the current study.

In the fully ligated 4T crystal, all strands are ligated in each direction. Its structural changes in response to tensile loading are presented in Figure 2a. After relaxation without any external forces, the crystal forms a rhombohedral shape rather than a cube, due to its motif design, with a tilt angle of ∼70° which is consistent with the experimental observation^23^. Upon initial loading in the vertical direction, the crystals begin to straighten with minimal resistance (Figure 2a(i)). As the applied force increases, the DNA edges in the crystal becomes straight and taut (Figure 2a(ii)). The cross-sectional area does not change significantly during this period. Further extension widens the gaps between the layers and base pairs start to dissociate. (Figure 2a(iii), inset). Accordingly, a gradual reduction is observed in the cross-sectional area. The connector (orange) strands also begin to be pulled toward the z-direction due to the ligation, suggesting a loss of the 3-fold rotational symmetry of the motif. At higher forces, most of the base pairs in the loading strands dissociate, and the structure transitions to ssDNA stretching (Figure 2(iv)). Areal change also accelerates in this stage, ultimately shrinking to nearly half of its original area, indicating a positive Poisson’s ratio. The simulation ends when the loaded unbound DNA eventually breaks.

**Figure 2.**
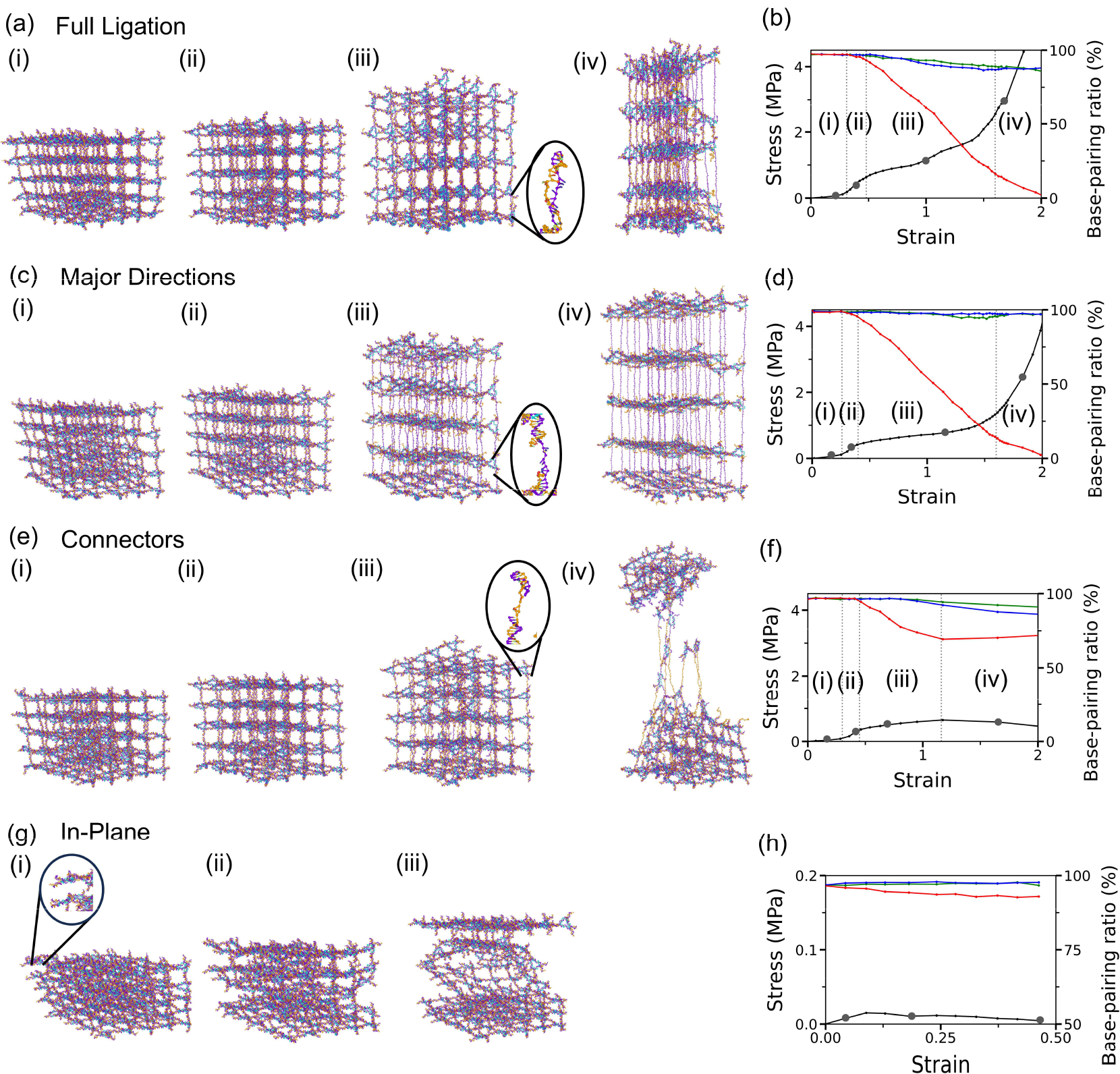
MD simulation results of 4T crystals with different ligation patterns. (a)-(b) Full ligation. (c)-(d) Ligation along major directions (x, y, and z axes). (e)-(f) Ligation with connectors. (g)-(h) In-plane ligation. For each ligation pattern, the configuration of the crystal with different deformation behavior under tensile loading are shown and corresponding data is marked by grey dots in stress vs. strain graph. In all plots, the base-pairing ratio of DNA duplexes in all three directions are plotted using the right axis. The directional color scheme is consistent with that in Fig. 1b with red representing the loading direction. The vertical dotted lines in the stress-strain plots indicate four distinct stages of deformation: (i) entropic elasticity, (ii) linear elasticity, (iii) dsDNA dissociation, and (iv) ssDNA stretching.

To further analyze this behavior, the stress vs strain curve is plotted in Figure 2b. Stress is calculated by dividing the applied force by the real time cross-sectional area, while strain is determined as the extension relative to the initial height. The force vs extension curve is provided in Supplementary Figure S1 and Table S2. The stress-strain plot reveals four distinct deformation stages. In stage (i), the structure is loose, and a small amount of stress (less than 0.2 MPa) leads to a large increase in strain (∼0.3). In stage (ii), both stress and strain increase linearly up to the yield point (∼0.7 MPa) indicating classical elastic behavior. Stage (iii) features a moderate increase in stress reaching ∼2.6 MPa as the structure extends until a strain of ∼1.6. This signifies that the crystal is extended more than 150%, resulting in a height twice its original value. Finally, in stage (iv), stress rises sharply until the strand breakage occurs. The data points associated with the configurations in Figure 2a are marked with grey dots on the plot.

To understand the deformation stages of fully ligated crystals, the base pairing ratio in all three directions is also plotted in Figure 2b using the right axis. The color of each curve corresponds to the directions defined in Figure 1c, where the red indicates z-direction — the direction of the applied tensile force. During stage (i), the base pairing is nearly intact in all directions with no significant dissociation. In stage (ii), a slight decrease in the red direction is observed, but it remains above 95%, suggesting minimal dissociation. In stage (iii), the crystal extends gradually with loading and the base-pairs along the loading direction drop significantly, falling below 15% eventually, which is defined as the end of the stage. By stage (iv), nearly all base pairs along the loading direction are dissociated with the ratio signifying a transition into ssDNA stretching. Meanwhile, the green and blue directions maintain base pairing ratios above 85% throughout the deformation process, indicating minimal impact on non-loading axes. Based on these results, four deformation stages are labeled as: (i) entropic elasticity, (ii) linear elasticity, (iii) dsDNA dissociation, and (iv) ssDNA stretching.

To examine the effect of ligation, a 4T crystal ligated along the major directions (i.e., purple strands) was simulated following the same procedure as the fully ligated case. As shown in Figure 2c(i), the initial structure appears slanted which begins to straighten under minimal tensile force. As the load increases, crystal extends proportionally, and DNA helices become taut (Figure 3c(ii)). Further stretching leads to gradual dissociation of base-pairs along the loading direction and increases the distance between adjacent planes (Figure 3c(iii)). With high forces, most base pairs are separated resulting in ssDNA stretching (Figure 3d(iv)) and the simulation ends upon strand breakage. Unlike the fully ligated crystal, the major direction ligation does not exhibit a significant reduction in the cross-sectional area during deformation. The corresponding stress and strain curve is plotted in Figure 2d. Like the fully ligated case, the plot can be divided into four deformation stages. The entropic (stage (i)) and linear elasticity (stage (ii)) extends structure with a stress of ∼0.4 MPa. Then, it enters stage (iii) or dsDNA dissociation regime with stress increasing up to 1.4 MPa. In stage (iv), stress rises sharply beyond 2 MPa due to the ssDNA stretching. The base-pairing ratio in three directions plotted in Figure 2d using the right axis provides further insight. During stage (i) and (ii), base-pairing along the loading direction remains nearly unchanged, indicating negligible dissociation. The ratio declines significantly in stage (iii), falling below 15%, and reaches 1% in stage (iv), confirming ssDNA stretching. The non-loading directions maintain base-pairing ratio above 95% throughout the simulation, consistent with the absence of cross-sectional area reduction. This suggests that the non-ligated orange strands connecting the planes are dissociating rather than being pulled with the loading strands. It is noted that the major direction ligation results in the yield stress approximately half that of the fully ligated crystal and requires less force to dissociate base pairs and reach stage (iv).

**Figure 3.**
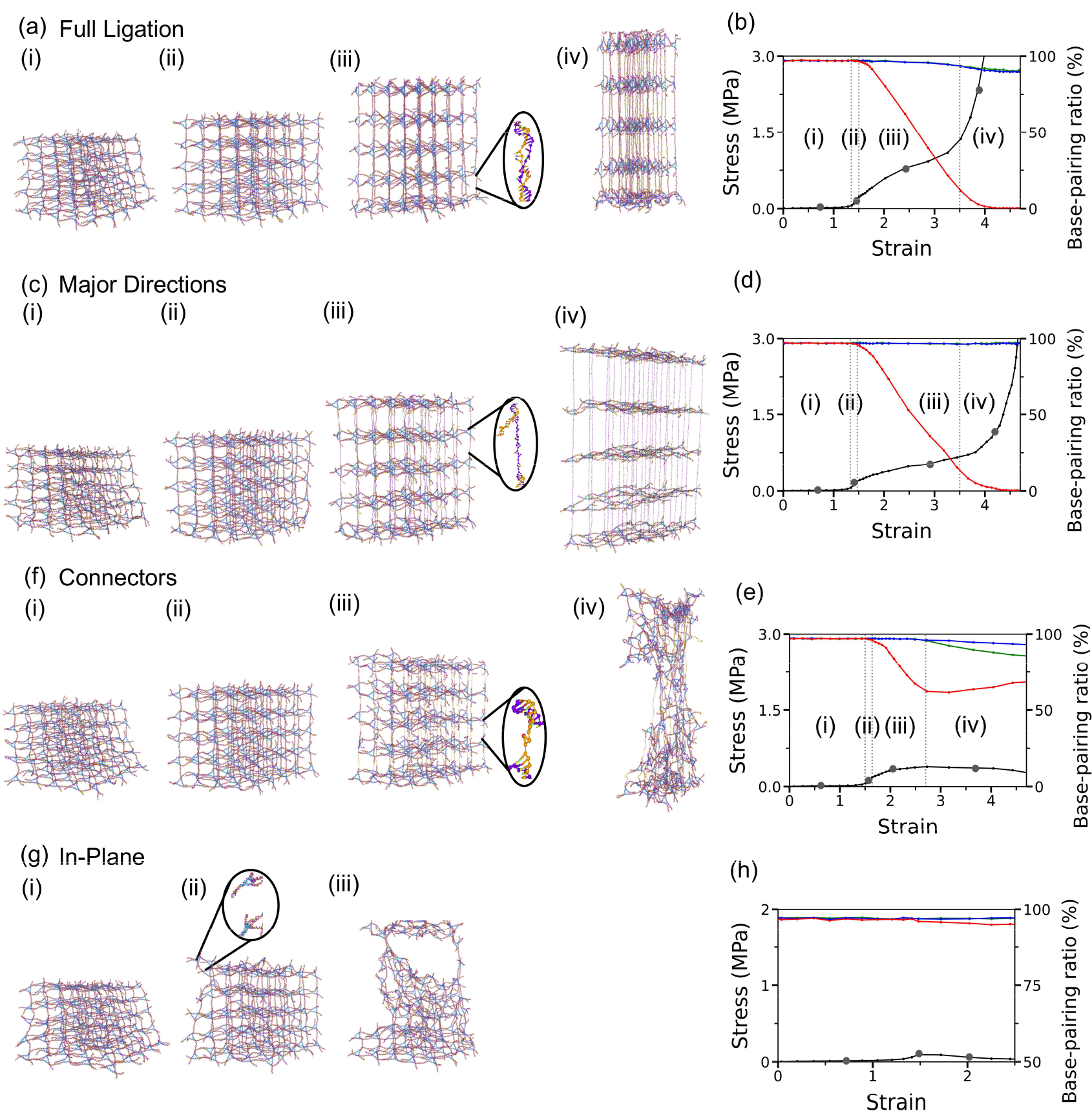
OxDNA results of 8T crystals with different ligation patterns, showing conformations under tensile loading and stress vs strain plots. The data points corresponding to the configurations are marked with grey dots on the plots. (a)-(b) Full ligation. (c)-(d) Ligation along the major directions. (e)-(f) Ligation with connectors. (g)-(f) In-plane ligation. The four deformation stages are also observed: (i) entropic elasticity, (ii) linear elasticity, (iii) dsDNA dissociation, and ssDNA stretching.

Next, we studied the connector ligated crystal, where the loading strands are not ligated. The relaxed structure has tilted DNA edges that are gradually straightened during the initial loading (Figure 2e(i)-(ii)). Upon further loading, however, the planes separate rather than stretch (Figure 2e(iii)) and the crystal fractures into two parts connected only by the orange strands (Figure 2e(iv)). Figure 2f presents the associated stress vs strain curve and base-pairing plots. The connector ligation displays (i) entropic elasticity regime up to a strain of ∼0.3, followed by (ii) linear elasticity region until reaching a yield strength of approximately 0.4 MPa. In stage (iii), the stress increases slowly with strain until reaching a critical point (∼0.6 MPa) where the structure breaks. During stage (iv), the stress decreases as the structure extends, ultimately leading to complete failure. Interestingly, the base association ratio along the loading direction shows a sharp decline in stage but begins to rise again slightly during stage (iv), suggesting that some sticky ends rehybridize after separation. Like other two ligation patterns, the connector ligated crystal also exhibits the four deformation stages with a similar slope in stage (ii). However, it undergoes different deformation behaviors where the crystal fails early due to the lack of ligation in the loading strands.

The final ligation pattern tested was in-plane ligation, in which neither the purple nor orange strands in the loading directions were ligated. In this case, the tilted structure does not straighten with minimal loading; instead, significant dissociation occurs at the sticky ends, leading to layer separation (Figure 2g(i)). Continued loading separates the planes even further and causes total structural failure (Figure 2g(ii)-(iii)). The stress-strain curve in Figure 2h confirms that in-plane ligation does not exhibit any distinct deformation stages. Instead, the applied stress remains low, indicating an easy failure. The base-pairing ratio in the x and y directions (green and blue) remains above 96%, while the red direction decreases slightly. This suggests that failure occurs layer by layer under applied tensile forces.

When comparing various ligation patterns, except for the in-plane ligation, the full ligation, major direction, and connectors result in a Young’s modulus ranging from 2 to 3 MPa (Figure S2; also see Supplementary Table S2 for stress and strain values at the end of each stage). After the linear stage, however, each ligation pattern exhibits unique deformation behavior under tensile loading. The fully ligated crystal exhibits the highest stress to achieve the same extension due to the increased resistance provided by additional ligated nucleotides. It is followed by the major directions and then the connectors. While both major directions and connectors involve the same number of ligation sites (half the total strands), they exhibit starkly different deformation behaviors. The connector ligated crystal fails during stage (iii) with a decrease in stress due to the absence of ligation in the loading strands. In contrast, the major direction case has continuously increasing stress throughout the simulation. The in-plane ligated crystal has more ligation sites than the major directions or the connectors, yet it breaks easily during stage (i). This is due to the lack of structural toughness arising from the absence of ligation along the loading direction (both purple and orange strands). The four ligation patterns have the same 4T motifs, but different number of ligation sites or locations lead to distinct and different deformation behaviors.

### MD Simulations of 8T crystals

The tensegrity triangle motif can be constructed with different edge lengths. To observe the effect of edge length on ligated crystals, we simulated 8T crystals with the same four different ligation patterns (Figures 3, S3, and S4 as well as Supplementary Table S3). The fully ligated structure is initially loose and relaxed (Figure 3a(i)). Compared to the 4T crystal, the 8T domain exhibits more tiled DNA edges with notable curvatures, due to a lower crossover density and larger cavity size, which results in greater structural flexibility. With increasing tensile forces, the initially relaxed strands along the loading direction become taut and the tilted structure becomes upright (Figure 3a(ii)). It is noted that the non-loading direction strands still exhibit some curvatures while no significant decrease in cross-sectional area is observed. Further extension increases the spacing between adjacent planes, with gradual base dissociation and a reduction in cross sectional area (Figure 3a(iii)). At higher forces, most base pairs in the loading strands are dissociated and the cross-section shrinks significantly (Figure 3a(iv)). Beyond this point, the crystal extends only slightly further before breaking and the end of the simulation. The corresponding stress vs strain curve in Figure 3b shows that the fully ligated 8T crystal has four deformation stages as well. In stage (i), during entropic elasticity region, the structure experiences a strain increase of approximately 1.4 under stress of ∼0.1 MPa. Stage (ii) shows linear elastic behavior up to a yield stress of ∼0.3 MPa. In stage (iii), the crystal is stretched further reaching a strain of about 3.5 during the stress increase up to ∼1.4 MPa. During stage (iv), the stress increases sharply, reaching a strain of 4 before failure. The base-pairing ratio reflects this progression where during stage (i) and (ii), the ratio is nearly constant with no significant decrease. In stage (iii), it gradually declines, indicating that base-pair dissociation occurs and leads to ssDNA stretching. Overall, the fully ligated 8T crystal forms more relaxed structure, and exhibits significantly longer entropic elasticity and dsDNA dissociation regions than 4T crystal due to its longer edge length.

The major direction (purple strands) ligated crystal also demonstrates unique patterns. Initially, the structure contains loose edges with some curvature (Figure 3b(i)) which straighten with the external loads (Figure 3b(ii)). With further extension, some base pairs dissociate (Figure 3b(iii)) and the loading strands eventually become unhybridized. Unlike the fully ligated crystal, the cross-sectional area remains nearly constant throughout the deformation process. The stress-strain curve and base-pairing ratios confirm this behavior (Figure 3d). In stage (i) and (ii), the structure reaches a strain of approximately 1.5 while stress was under ∼0.2 MPa. Then, stress increases by 0.4 MPa only, while strain increases from about 1.5 to 3.5. Due to the absence of ligation on half of the strands, the stress is lower compared to the fully ligated crystal. In the final stage (iv), stress increases exponentially, exceeding 3 MPa, and the structure fails. The base-pairing along the loading direction drops significantly in stage (iii) and reaches ∼1% in stage (iv). Note that the base association ratio in the non-loading directions remains unchanged, consistent with the observation that the cross-sectional area does not change throughout deformation.

The connector ligated 8T crystal (ligation on the orange strands) exhibits a similar initial trend as the above two cases. As tensile forces are applied, the crystal shows entropic and linear elasticity behaviors (Figure 3e(i)-(ii)). Continued loading leads to the dissociation of base pairs in the loading direction. (Figure 3e(iii)). Shortly after, the crystal fractures, splitting into two parts that remain connected by ligated orange strands. The stress-strain curve in Figure 3f reveals that the combined entropic and linear elasticity reaches a strain of ∼1.6. In stage (iii), the stress begins to decrease at strain of ∼2.7, indicating a crystal failure. Further extension completely breaks the structure as shown in Figure 3e(iv). Like the behavior of the 4T connector ligation, the base-pairing ratio decreases significantly during the dsDNA dissociation, and it recovers slightly in stage (iv), indicating partial rehybridization of the sticky-ends after separation.

Finally, the in-plane ligation demonstrates a very weak crystal structure against tensile loading. Upon pulling in the orthogonal direction, the structure does not elongate; instead, it breaks apart layer by layer. The stress vs strain curve in Figure 3h indicates no distinct deformation stages. The crystal shows an entropic elasticity behavior region at minimal stress, but a linear elasticity regime is not observed due to failure. The base association ratios in all three directions remain nearly constant.

### Effect of Motif Length on the Mechanics of DNA Crystals

From the simulations of 4T and 8T crystals, unique behaviors are observed across different ligation patterns. One of the primary distinctions is found in the entropic elasticity regime. Crystals with longer motif lengths exhibit more relaxed structures, requiring a prolonged straightening process. To further investigate this trend, four fully ligated crystals with varying motif lengths (2T,4T, 6T, and 8T) are compared with 2T and 6T results presented in Supplementary Figures S5 and S6. Figure 4 compares the deformation behaviors of individual DNA helices in each crystal. For consistency, the applied plane forces are divided by 25 to approximate the force acting on an individual duplex. The curves are shifted horizontally to align the linear elasticity region for a better comparison. As the motif length increases from 2T to 8T, the crystals exhibit progressively longer entropic elasticity period. The 2T crystal displays no noticeable entropic elasticity, whereas the 4T, 6T, and 8T structures show strain values of approximately 0.3, 1, and 1.4, respectively. Despite these differences, all crystals exhibited nearly identical slopes in the linear region and shared a consistent yield point at around 80 pN.

**Figure 4.**
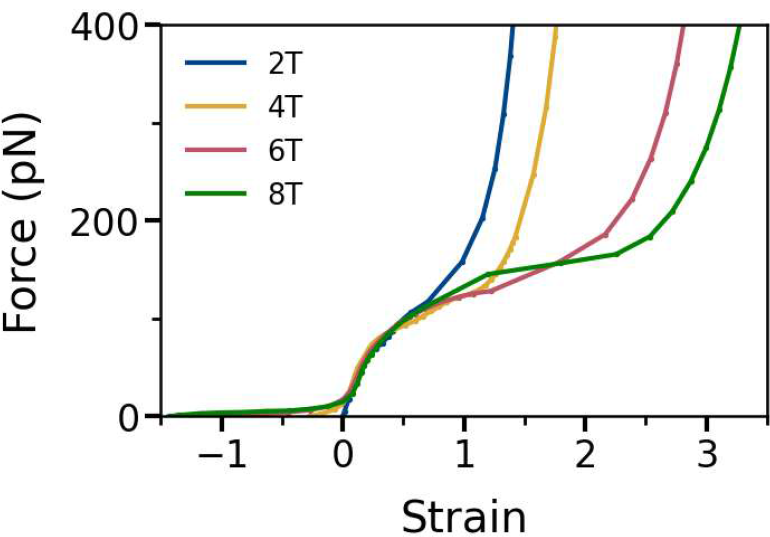
Force vs. strain plots for each DNA duplex in the fully ligated crystals with various motif lengths. The linear elasticity stage are shifted for a better comparison. As the motif length (number of helical turns) increases, the entropic elasticity regime also increases. The 2T crystal exhibits no entropic stage, while the 8T case has the longest. The similar trend is also reflected in strain range, as crystals with shorter motifs (2T) have less nuelceotides in the structure, the overall extension is shorter and greater forces are required for a given strain compared to other cases.

Motif length also influences the overall extension of the structures significantly. Longer motifs, which contain more nucleotides in the loading strands, require greater extension before complete base-pair dissociation and transition into ssDNA stretching. For instance, ssDNA stretching begins at strain of approximately 0.7 and 1 for the 2T and 4T crystals, respectively. The 6T and 8T crystals transition at higher strains, with complete dissociation occurring around strain values of ∼2 and 2.5. As a result, the total extension (excluding entropic elasticity) reaches a strain of up to about 3 for the 8T crystal (a 300% increase in height), while it is limited to ∼1 for the 2T case. This behavior is similar to pulling a single duplex of varying length where shorter strands reach structural failure with less extension, whereas longer strands can endure greater elongation. These findings provide useful information on how motif length influences structural flexibility and extension, which is crucial in designing dynamic and deformable DNA structures.

## CONCLUSIONS

In this study, we investigated the effect of ligation patterns and motif lengths on mechanical behavior of DNA crystals using coarse-grained MD simulation. We observed that ligation patterns influence the mechanical strength and failure behavior of DNA crystals. Fully ligated structures have the highest yield point and structural integrity, whereas the crystals ligated along the major directions require less force to reach the same extension. Other ligation patterns, where the loading directions are not ligated, result in significantly weaker elastic behaviors and early structure failure. Besides the ligation patterns, the motif length also greatly impacts structural rigidity and extension in response to mechanical loading.

This work provides essential insight into structural mechanics of DNA crystals, which may be extended for designing DNA assemblies as it allows researchers to predict how the structures will response under various conditions. Depending on whether a flexible, tightly packed, highly stretchable, or fragile structure is desired, our results can recommend an optimal motif design for task-specific functions. For instance, long motif lengths result in large cavity sizes, which can be utilized to encapsulate proteins or nanoparticles for biosensing applications. On the other hand, short motifs like 2T crystal are suitable for sturdy structures. By modifying ligation patterns, it is also possible to selectively weaken strand connections and potentially control dissociation at specific sites with an external force application. For example, in-plane ligation may be used to separate 2D layers by applying external loads along the weak direction. Furthermore, DNA functionalization with inorganic materials has been actively studied, using DNA crystals as a template for creating 3D architected inorganic materials. Silicification can enhance mechanical stiffness^44^, while coating with metals or semiconductors can alter electrical and optical behaviors. If inorganic functionalization is performed on in-plane ligated crystals, it can potentially produce 2D layers of inorganic crystal via micromechanical exfoliation, as with graphene or other 2D materials such as MoS^2^ and WSe^2^ flakes. In this work, only four simple ligation patterns were examined, yet more complex designs such as two different ligation patterns in a single crystal could exhibit interesting behaviors and further expand the utility of DNA crystals in nanotechnology.

## ACKNOWLEDGMENTS

This work was supported by the U.S. Department of Energy (DOE), Office of Science, Basic Energy Sciences (BES) under award no. DE-SC0020673. C.M. acknowledges the support from the U.S. National Science Foundation (NSF) under award no. 2025187.

