## Supplementary Information for "Understanding Structural Mechanics of Ligated DNA Crystals via Molecular Dynamics Simulation"

| | Cyan | Purple | Orange |
|---|---|---|---|
| 2T | ACACCGTACACCGTACACCGT | GAGCAGCCTGTACGGACATCA | TCTGATGTGGCTGC |
| 4T | CGGTATTCACCAGGATGCGGTA<br>TTCACCACGATGCGGTATTCAC<br>CACGATG | GAAAAACACTGCCTGAATACCG<br>CATCGTGGACTGACTCAAAA | TCTTTTGAGTCAGTG<br>GCAGTGTTTT |
| 6T | ATGCGGTCAGTAATTCACCACG<br>AGCTAGATGCGGTCAGTAATTC<br>ACCACCAGCTAGATGCGGTCAG<br>TAATTCACCACGAGCTAG | GAGTAACAGCATACTGCCTGAAT<br>TACTGACCGCATCTAGCTCGTG<br>GACTGATCGACCTCAGAG | TCCTCTGAGGTCGAT<br>CAGTGGCAGTATGCT<br>GTTAC |
| 8T | CGGTCAGTAATTCACCACGAGC<br>TAGATGCGGTCAGTAATTCACC<br>ACGAGCTAGATGCGGTCAGTAA<br>TTCACCACGAGCTAGATG | GAGGAGCAAACTTCTAACAGCAT<br>ACTGCCTGAATTACTGACCGCAT<br>CTAGCTCGTGGACTGATCGACC<br>TCCTTGAAAGACAGAG | CTCCTCTGTCTTTCAA<br>GGAGGTCGATCAGTG<br>GCAGTATGCTGTTAG<br>AAGTTTGCTC |

**Table S1.** The sequence information for the tensegrity triangle motif with different numbers of helical turns (2T, 4T, 6T, and 8T). Each sequence is listed from 5’ to 3’. Note 2T, 4T, and 6T motifs have 2-nt sticky ends, while 3-nt is used for the 8T crystal.

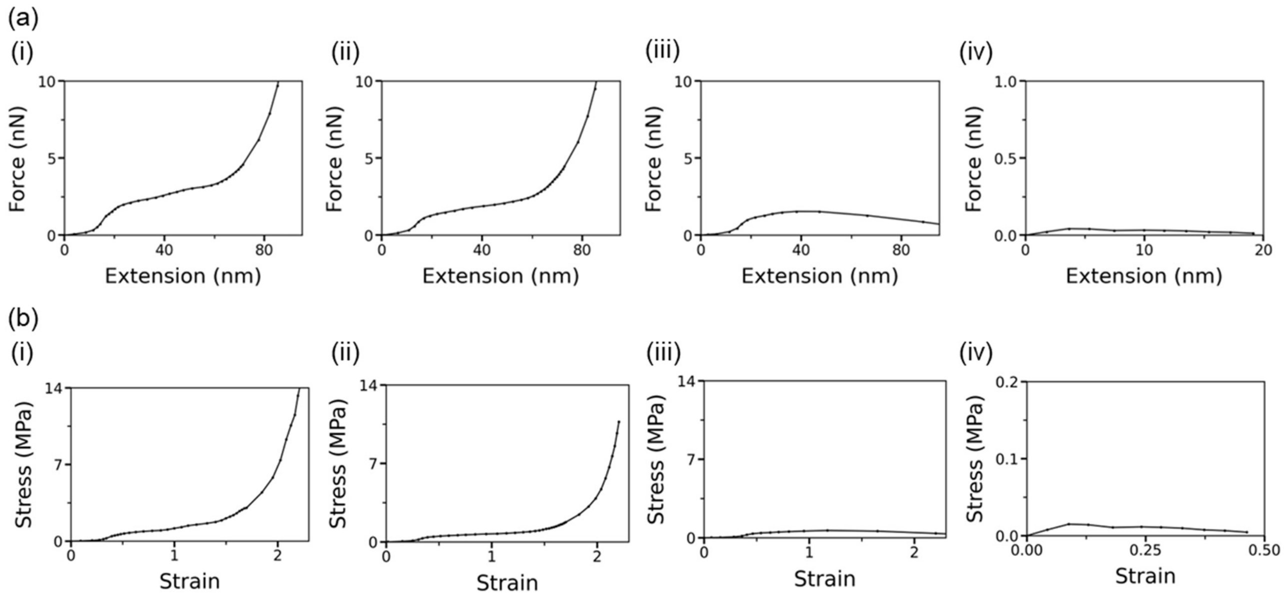


**Figure S1.** (a) Force vs extension and (b) stress vs strain plots for 4-turn crystals with different ligation patterns. (i) Full ligation, (ii) major directions, (iii) connectors, and (iv) in-plane ligation. The stress-strain curves in Figure 2 are plotted up to 4 MPa to highlight distinct deformation stages, while Figure S1(b) presents the data with extended y-axis to illustrate the overall deformation.

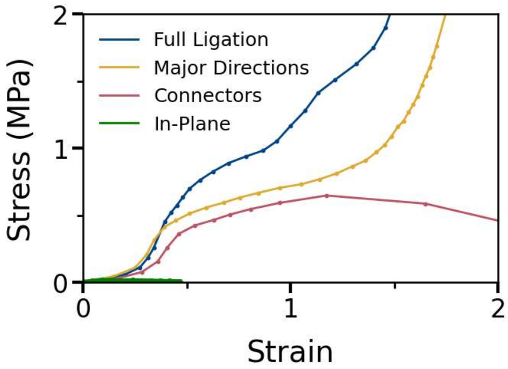


**Figure S2.** Stress vs strain curves for various ligation patterns of the 4T crystal. Full ligation as well as ligation with major directions and connectors exhibit similar slopes in the linear elasticity range, with a Young's modulus of approximately 2-3 MPa. However, the difference is shown in yield points. The full-ligation case has the highest yeid stress at ~0.7 MPa, while those with major directions and connectors show nearly a half value. In contrast, the in-plane ligation structure fails before entering the linear elastic regime. It is also notable that the crystals with full ligation and major directions increase sharply beyond strain > 1, whereas the case of ligation with connectors show a clear decrease. With unligated along the loading direction, the crystal with in-plane ligation fails even before entering the linear elasticity stage.

| | Entropic Elasticity | Linear Elasticity (Yield Stress) | dsDNA Dissociation |
|---|---|---|---|
| Full Ligation | ~ 0.2 MPa, 0.3 | ~ 0.7 MPa, 0.5 | ~ 2.6 MPa, 1.6 |
| Major Directions | ~ 0.2 MPa, 0.3 | ~ 0.4 MPa, 0.4 | ~ 1.4 MPa, 1.6 |
| Connectors | ~ 0.1 MPa, 0.3 | ~ 0.4 MPa, 0.5 | ~ 0.6 MPa, 1.2 |

**Table S2.** Stress and strain values of 4T crystals at the end of each deformation stages. In-Plane ligation is not included as it does show any stages.

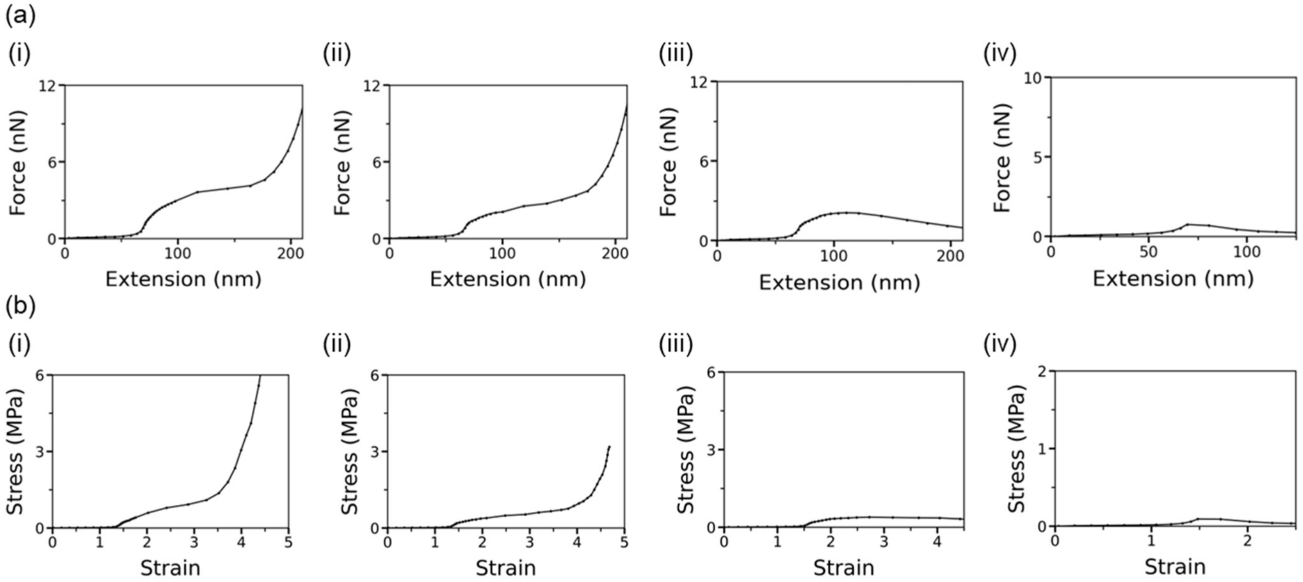


**Figure S3.** (a) Force vs. extension and (b) stress vs. strain curves with expanded y-axis for 8-turn crystals. (i) Full ligation, (ii) major directions, (iii) connectors, and (iv) in-plane ligation.

| | Entropic Elasticity | Linear Elasticity (Yield Stress) | dsDNA Dissociation |
|---|---|---|---|
| Full Ligation | ~ 0.06 MPa, 1.4 | ~ 0.3 MPa, 1.5 | ~ 1.4 MPa, 3.5 |
| Major Directions | ~ 0.06 MPa, 1.4 | ~ 0.2 MPa, 1.5 | ~ 0.6 MPa, 3.5 |
| Connectors | ~ 0.06 MPa, 1.5 | ~ 0.2 MPa, 1.6 | ~ 0.4 MPa, 2.7 |

**Table S3.** Comparison of stress and stain values for 8T crystals at the end of each deformation stages.

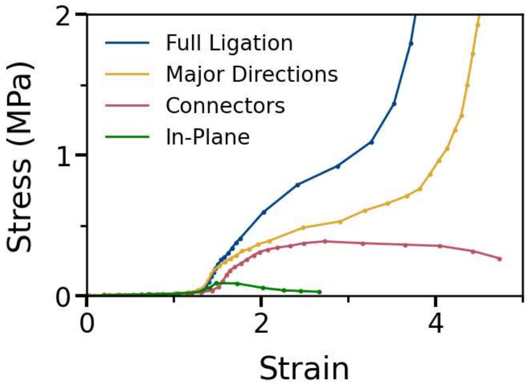


**Figure S4.** Stress vs strain curves for the 8T crystal with various ligation patterns. All cases have a similar strain range during the entropic elasticity period and a similar slope in linear elasticity range. Significant differences are observed after the linear range. The fully ligated crystal shows highest stress throughout the whole stage. The crystal with ligation along major directions has lower stress than fully ligated case but still shows a similar trend of the drastic increase in the ssDNA stretch regime. Connectors and in-plane ligation patterns exhibit a decrease beyond strain of around 3 and 1.5, respectively, due to the lack of ligation along the loading direction.

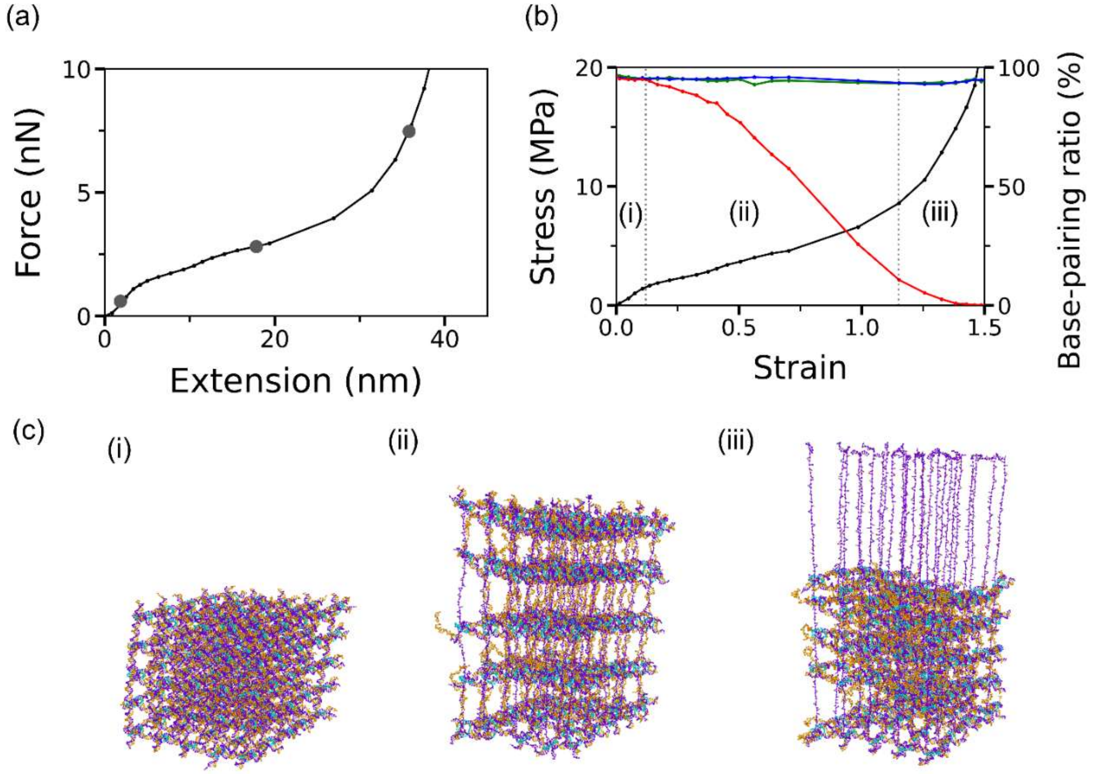


**Figure S5.** MD simulation results of a fully ligated 2T crystal (made of 5×5×5 motifs) under tensile forces. (a) Force vs. extension plot and (b) stress vs. strain curve along with dsDNA base pair association ratio (%) in three directions indicated by red, blue, and green. Stress is calculated by dividing a pair of plane forces by the cross-sectional area at each data point. Strain is computed by dividing the extension (elongation along the loading direction) by the initial height of the crystal. Three deformation stages are illustrated with vertical dotted lines: (i) linear elasticity, (ii) dsDNA dissociation, and (iii) ssDNA stretch. (c) The configurations of 2T crystal at different stages, corresponding to the points marked with grey dots in (a). Unlike other structures, the 2T crystal shows only three regimes. The linear elasticity range starts immediately upon tensile forces without the entropic elasticity period. The yield stress is recorded at about 1 MPa (~1.4 nN). Stage (ii) persists until the stress reaches about 8.5 MPa (~4.8 nN). In stage (iii), the planes no longer extend along the loading direction. Instead, they collapse and stack on top of each other. This deformation behavior is also reflected on the base pairing association ratio where the loading direction (red) decreases drastically in stage (ii), while those along the in-plane (blue and green) directions are not significantly affected.

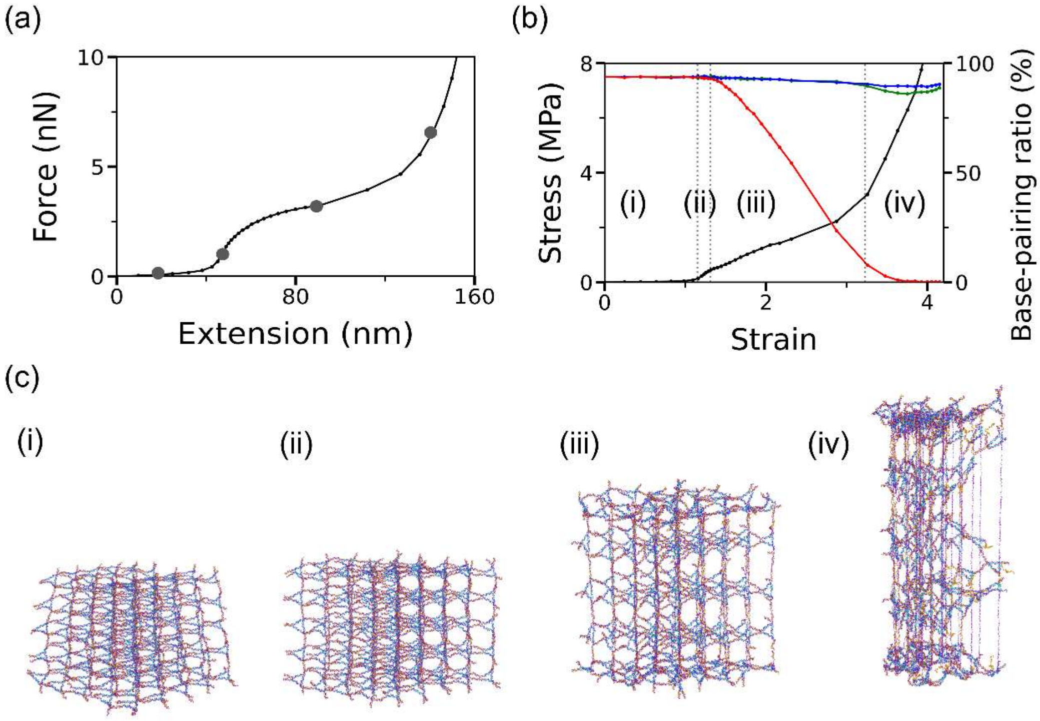


**Figure S6.** (a) Force vs. extension plot and (b) stress vs. strain curve of a fully ligated 6T DNA crystal under tensile loading. The plot also shows the dsDNA base pair ratios along the loading (red) and in-plane (blue, green) directions using the right axis. The MD computation shows four distinct stages, which are separated by vertical dotted lines: (i) entropic elasticity, (ii) linear elasticity, (iii) dsDNA dissociation, and (iv) ssDNA stretch. Compared to 2T and 4T crystals, it has a longer entropic elasticity stage (i) (strain of 1), but shorter than the 8T crystal. (c) The conformations corresponding to the data marked with grey circles in (a).
